# Single virus assay reveals membrane determinants and mechanistic features of Sendai virus binding

**DOI:** 10.1101/2021.11.23.469711

**Authors:** Amy Lam, Orville O. Kirkland, Papa Freduah Anderson, Nandini Seetharaman, Dragan Vujovic, Patricia A. Thibault, Kristopher D. Azarm, Benhur Lee, Robert J. Rawle

**Affiliations:** Dept of Chemistry, Williams College, Williamstown, MA; Dept of Microbiology, Icahn School of Medicine at Mount Sinai, New York, NY

**Author notes:** Address correspondence to Robert J. Rawle.

## Abstract

Sendai virus (SeV, formally murine respirovirus) is a membrane-enveloped, negative sense RNA virus in the *Paramyxoviridae* family, and is closely related to human parainfluenza viruses. SeV has long been utilized as a model paramyxovirus and has recently gained attention as a viral vector candidate for both laboratory and clinical applications. To infect host cells, SeV must first bind to sialic-acid glycolipid or glycoprotein receptors on the host cell surface via its hemagglutinin-neuraminidase (HN) protein. Receptor binding induces a conformational change in HN, which allosterically triggers the viral fusion (F) protein to catalyze membrane fusion. While it is known that SeV binds to α2,3-linked sialic acid receptors, and there has been some study into the chemical requirements of those receptors, key mechanistic features of SeV binding remain unknown, in part because traditional approaches often convolve binding and fusion. Here, we develop and employ a fluorescence microscopy-based assay to observe SeV binding to supported lipid bilayers (SLBs) at the single particle level, which easily disentangles binding from fusion. Using this assay, we investigate mechanistic questions of SeV binding. We identify chemical structural features of ganglioside receptors that influence viral binding and demonstrate that binding is cooperative with respect to receptor density. We measure the characteristic decay time of unbinding and provide evidence supporting a “rolling” mechanism of viral mobility following receptor binding. We also study the dependence of binding on target cholesterol concentration. Interestingly, we find that while SeV binding shows striking parallels in cooperative binding with a prior report of Influenza A virus, it does not demonstrate a similar sensitivity to cholesterol concentration and receptor nano-cluster formation.

**STATEMENT OF SIGNIFICANCE:** Paramyxoviruses are a family of membrane-enveloped viruses with many notable human and animal pathogens. In this study, we develop and use an assay to observe the initial step of infection – virus binding to the host membrane – for Sendai virus, the prototypical paramyxovirus, at the single virus level. This assay uses cell membrane mimics – supported lipid bilayers – as targets for virus binding to enable easy control of the membrane components with which the virus interacts. Using our assay, we gain insight into basic biophysical questions about Sendai virus binding, including the chemical characteristics of the receptor, the cooperative nature of binding, the influence of cholesterol, and the mechanism of viral mobility following binding.

## INTRODUCTION

Paramyxoviruses are a family of enveloped, non-segmented negative sense RNA viruses which include the human parainfluenza viruses, measles virus, Newcastle disease virus, various veterinary viruses, and are responsible for significant human and animal disease worldwide. Sendai virus (SeV), formally murine respirovirus, has been a prototypical and well-studied model virus of the *Paramyxoviridae* family, and is quite interesting in its own right (1, 2). It primarily causes disease in rodents and other animals, and while it can infect human cells, it does not cause disease in humans or lead to oncogenic transformation (2, 3). As such, it has been developed, among other uses, as a gene therapy vector (2), as a CRISPR/Cas9 delivery system (3), as a commercial vector (ThermoFisher) to induce pluripotent stem cells (4), as well as a vaccine candidate backbone for HIV (5), RSV (6), SARS-CoV-2 (7), and others (8).

For all enveloped viruses, the first step in infection is binding of the virus to a receptor or attachment factor on the host cell membrane surface. SeV, as with other closely related paramyxoviruses, utilizes α2,3-linked sialic acid glycolipids or glycoproteins as receptors for cell entry (9–12). Receptor binding is mediated by the hemagglutinin-neuraminidase (HN) receptor binding protein (formerly called the HN attachment protein), a transmembrane protein embedded in the viral envelope. HN is commonly understood to be arranged in a dimer of dimers, and composed of a helical stalk domain with a globular head which contains the receptor binding pocket (13, 14). Upon receptor binding, a conformational change is induced in HN, which then allosterically triggers the viral F protein to catalyze membrane fusion with the host cell plasma membrane, initiating infection (13).

While it is known that SeV utilizes α2,3-linked sialic acid receptors for cell entry, and there has been some study into the chemical requirements of those receptors in non-membrane systems (9, 10, 12, 15, 16), key features of SeV binding and behavior following binding remain unknown, including the cooperativity of viral binding, the influence of receptor localization in the target membrane, and viral mobility following receptor binding.

In this report, we examine these key features of SeV binding using a fluorescence microscopy-based single virus binding assay. Single virus assays have been employed fruitfully for other families of viruses to study binding and membrane fusion (17–33). These assays are commonly implemented using synthetic target membranes such as supported lipid bilayers (SLB) or tethered liposomes which afford the researcher precise control over membrane topology, composition, and environment to ask specific biophysical questions. To our knowledge, such assays have not been employed for any paramyxovirus. Here, we present the development and validation of a single virus binding assay for SeV. We then utilize this assay to ask targeted questions about viral binding and viral behavior following binding. We investigate the role of sialic acid density and positioning for receptor binding, the cooperative dependence of binding on receptor density, the influence of cholesterol composition, and the mechanism of viral mobility following binding.

## RESULTS AND DISCUSSION

### Assay overview and development

Our single virus binding assay for SeV (Figure 1A) was adapted from a previously reported influenza A virus (IAV) assay (23), but with some important modifications and considerations. Viral particles were fluorescently labeled with a lipophilic dye and were introduced at a known concentration (typically 0.11 nM, see Supporting Information for concentration determination) into a polydimethylsiloxane (PDMS) microfluidic chamber housing a glass-supported bilayer. The SLB was prepared with one of several sialic acid glycolipids, which served as receptors for viral binding. Following incubation, unbound viruses were removed via buffer rinse, and bound viruses were then imaged by fluorescence microscopy.

**Figure 1.**
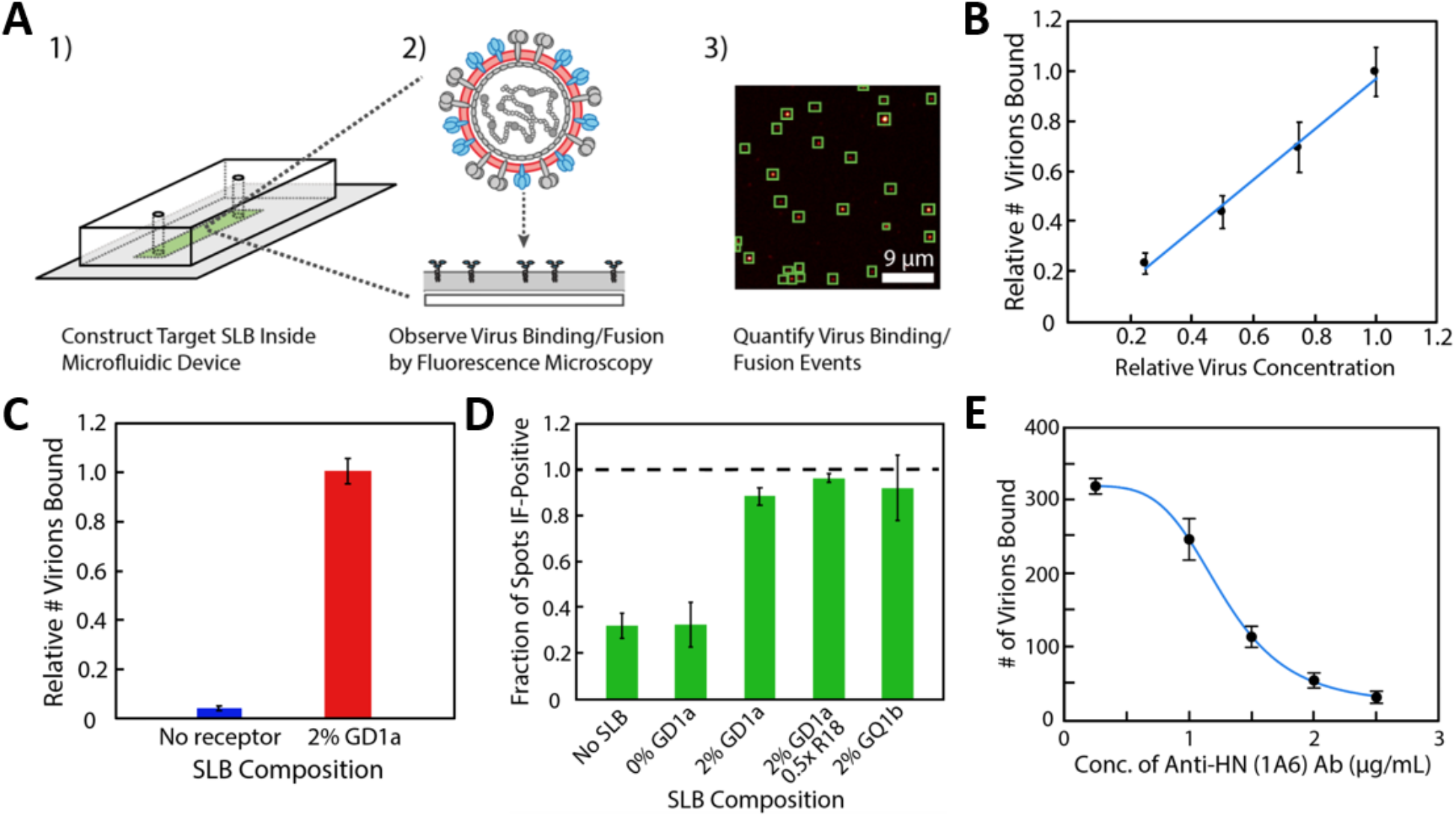
Overview of single virus binding assay design and validation data. A) shows a schematic of the assay design. Supported lipid bilayers (SLBs) are self-assembled inside a microfluidic device. Sendai virions, membrane-labeled with Texas Red-DHPE (TR) or R18, are introduced into the flow cell, where they can bind to receptors in the SLB. Binding and, in limited cases, fusion are observed and quantified by fluorescence microscopy. B) shows a linear relationship between the viral concentration added to the flow cell, and the relative number of virions bound. In these measurements, the SLB contained 2% GD1a receptor and the viral concentration ranged from 0.025 nM to 0.1 nM. Error bars are ± standard error of ≥ 3 sample replicates, and ≥ 12 separate image locations within each sample. C) depicts the relative number of virions bound to SLBs containing either 2% GD1a or no receptor. Very little binding is observed to SLBs without receptor. Error bars are ± standard error of ≥ 3 sample replicates, and ≥ 10 separate image locations within each sample. D) shows the fraction of spots bound to various SLBs which show positive immunofluorescence (IF) labeling by anti-HN (1A6) antibody, followed by Alexa-488-labeled secondary antibody. Colocalization between Alexa-488 and the membrane label in the particles (TR or R18) was used to determine if a particle was IF-positive. “No SLB” indicates that particles were attached non-specifically to the glass coverslip instead of an SLB, followed by surface passivation by 30 g/L bovine serum albumin. “0.5x R18” indicates that viral particles were labeled with R18 at 0.5X concentration (see Materials and Methods for labeling details); all other data in this panel was collected with TR-labeled particles. Error bars are ± standard error of ≥ 3 sample replicates with ≥ 10 separate image locations within each sample; the “0.5x R18” sample had 1 sample replicate and error shown is standard deviation of 10 separate image locations. E) shows the sensitivity of Sendai virus binding to 2% GD1a SLBs following pre-treatment of the virus with 1A6 antibody. Virus labeled with R18 (1x) was incubated with 1A6 antibody at the concentrations shown for 30 minutes on ice prior to injection into the flow cell. The blue line shows the best fit to a sigmoid curve; IC_50_ = 1.3 ± 0.1 µg/mL. Error bars are ± standard deviation of 6-10 separate image locations within each sample.

We found that three critical aspects of the assay design were 1) the method of introduction of the virus into the microfluidic chamber, 2) the extent of labeling and identity of the fluorescent dye incorporated into the viral envelope, and 3) the minimization of receptor-triggered membrane fusion by temperature control. Some of these issues are particular to paramyxoviruses and similar viruses, such as the minimization of fusion triggered by receptor binding, but others are relevant to viral binding and fusion studies more broadly. If not carefully controlled, each can lead to unwanted artifacts (such as non-specific binding or fusion) and/or difficulties in data collection; this is discussed in detail in the Supporting Information (see Figures S2-S6 and associated text). We highlight that proper fluorescence labeling is an issue of particular note, as many previous reports studying SeV fusion have utilized fluorescent labeling of the viral envelope (see for example refs (34–38)). Depending on the extent of labeling, non-specific fusion artifacts may have been occurring.

### Assay validation

After optimizing the issues identified above, we performed various validation experiments. First, we observed that viral binding to SLBs with sialic acid receptor (2 mol% GD1a) was dose-dependent with concentration of virus added (Figure 1B). Second, we verified that SeV showed little binding to SLBs without receptor, as compared to SLBs with 2% GD1a (Figure 1C). Third, using immunofluorescence (IF) imaging of HN proteins on stably bound particles, we observed a much higher percentage of IF-positive particles bound to SLBs with 2 mol% GD1a (88.5% ± 3.8%) compared to particles bound to SLBs without receptor (32.3% ± 9.9%) or bound non-specifically to a glass surface (31.9% ± 5.5%), (Figure 1D). Control experiments using secondary antibody only showed negligible IF-positive results (Figure S1). This indicates that receptor binding to SLBs efficiently selects binding-active virions from non-viral or quasi-viral particles which are co-purified, a recently identified challenge in viral purification (39–41), and a potential yet often ignored confounding factor in single virus experiments (See Supporting Information for further discussion).

Finally, to verify that specific HN-receptor interactions were responsible for viral binding in our assay, we examined antibody inhibition of SeV binding by pre-treating the virus with a neutralizing anti-HN monoclonal antibody (1A6) prior to introduction into the flow cell. We determined the IC_50_ value in our assay at 1.3 ± 0.1 µg/mL (Figure 1E) and at saturating antibody concentrations we observed a ∼10 fold decrease in binding to SLBs with 2% GD1a compared to virus which had been mock treated with buffer only. This antibody inhibition was observed both with TR-labeled virus and R18-labeled virus (at appropriate R18 concentrations) indicating a robust antibody response in our assay independent of dye identity (Figure S7). Conversely, no inhibition of apparent binding was observed to SLBs without GD1a, supporting the IF results above indicating that particles bound to membranes without receptor are largely bound non-specifically or are non-viral particles. Similar inhibition was also observed with a separate neutralizing anti-HN antibody (3G12), see Figure S8. Interestingly, we also observed moderate inhibition of binding with an anti-F monoclonal antibody (11H12) (42), see Figure S8. This suggests that F and HN proteins may be closely intermixed on the viral surface, such that antibody binding to F leads to partial inhibition of HN receptor binding by steric hindrance. Together, these antibody inhibition results underscore that HN-receptor interactions are responsible for viral binding in our assay. More broadly, they also demonstrate the capability of assays such as these to cleanly monitor antibody inhibition of binding to specific receptors without convolving with other steps in the infectious cycle such as membrane fusion.

### Characterization of viral behavior and studies of viral binding mechanisms

To our knowledge, single virus measurements of this type have not been conducted for SeV or other paramyxoviruses. Therefore, we performed various experiments/measurements to characterize viral behavior and to examine specific questions relating to viral binding, including the influence of receptor chemical structure, cooperativity of binding, dependence on cholesterol concentration, and mechanism of mobility following binding. These are each described sequentially below.

### Number and positioning of α2,3-linked sialic acids on receptors directly modulates viral binding

Number, positioning, and chemical linkage of sialic acid (SA) receptors can be an important modulator of viral attachment for SA-binding viruses, and can play a role both in host and tissue tropism (43). Previous work has indicated that SeV and other SA-binding paramyxoviruses such as mumps virus and HPIV3 have a preference for α2,3-linked SA residues on either glycolipids or glycoproteins (13). To study the influence of the number and positioning of α2,3-linked SA residues on SeV binding in our assay, we studied binding to membranes containing 2 mol% of different SA-gangliosides: GM3, GM1, GD1a, or GQ1b (Figure 2). GM1, GD1a, and GQ1b all possess the same ganglio-tetrose backbone with 1, 2, or 4 SA residues either branched internally off the 2^nd^ backbone sugar and/or in the terminal position as depicted (Figure 2A). GM3 is similar in structure to GM1, with 1 SA residue, but is lacking the final 2 sugars of the tetrose backbone, which transforms its SA into a terminal residue rather than an internal branching residue.

**Figure 2.**
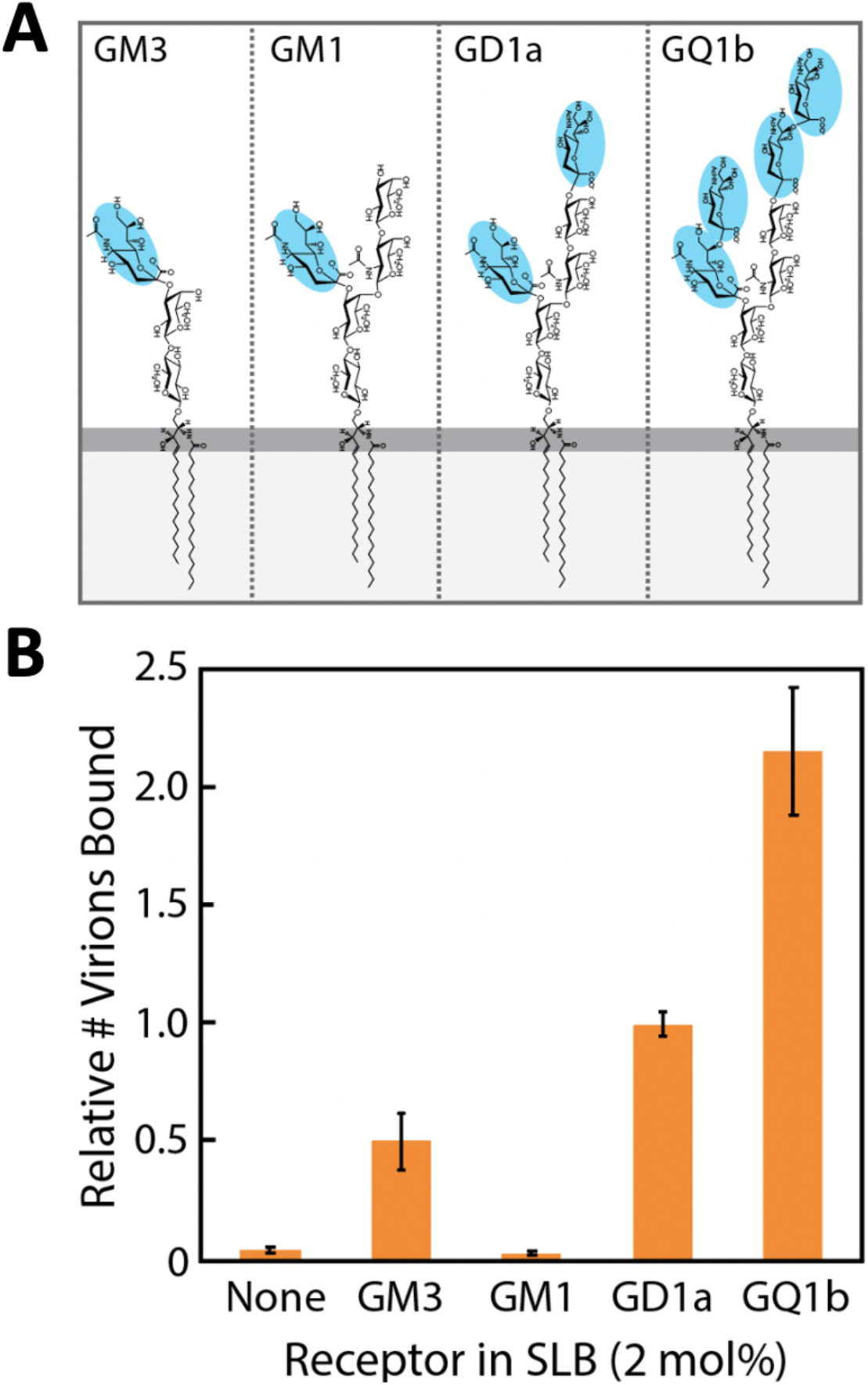
Chemical structure of gangliosides in SLBs directly modulates SeV binding. Single virus binding measurements were performed to SLBs containing 2% GM3, GM1, GD1a, GQ1b, or no receptor. A) shows the chemical structure of each receptor used. Sialic acids are highlighted in blue. Note that GM1, GD1a, and GQ1b each possess the same ganglio-tetrose backbone, whereas GM3 lacks the final 2 sugars of the tetrose backbone. B) shows the relative number of virions bound to SLBs with the different ganglioside receptors. Binding measurements are shown relative to 2% GD1a. Error bars are ± standard error of ≥ 4 sample replicates, and ≥ 10 separate image locations within each sample.

We observed that while binding to 2% GM1 membranes occurred at nearly identical levels to membranes containing no receptor at all, membranes containing 2% GM3 elicited a moderate amount of viral binding. This indicates that the final 2 sugars of the tetrose backbone in GM1 likely inhibit access to the internally branched SA, preventing effective binding by HN. On the other hand, GD1a, with an identical internally branched SA as GM1, but also containing an additional terminal SA, exhibited approximately 2X greater viral binding than GM3. This suggests that either the terminal SA of GD1a is a much more favorable binding target for HN, or that the addition of the terminal SA alters the ganglioside head group structure such that the internally branched SA becomes more accessible than in GM1. GQ1b exhibited the highest level of binding, approximately 2X greater than GD1a, suggesting that the additional sialic acid residues (4 total for GQ1b), as well as the greater distance of the terminal sialic acids from the tetrose core facilitate increased binding by HN (Figure 2B).

### Sendai virus exhibits different cooperative binding to different ganglioside receptors

Monomeric receptor binding affinities of HN for the closely related HPIVs are rather weak (*K*_*d*_ ∼0.1mM), (44) suggesting that avidity rather than affinity likely governs SeV binding, and opens the possibility that binding may be cooperative. To directly study the cooperativity of SeV binding, we observed virus binding to SLBs with varying GD1a concentrations (Figure 3A). We observed a characteristic sigmoidal response to GD1a concentration, indicative of cooperativity. To quantify the extent of cooperativity, we fit the data to a Hill model binding curve:

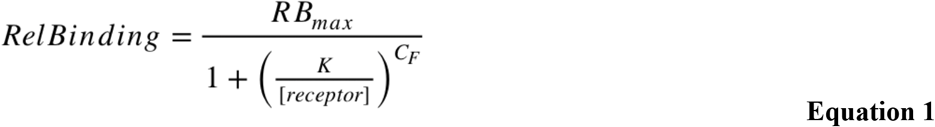

**Figure 3.**
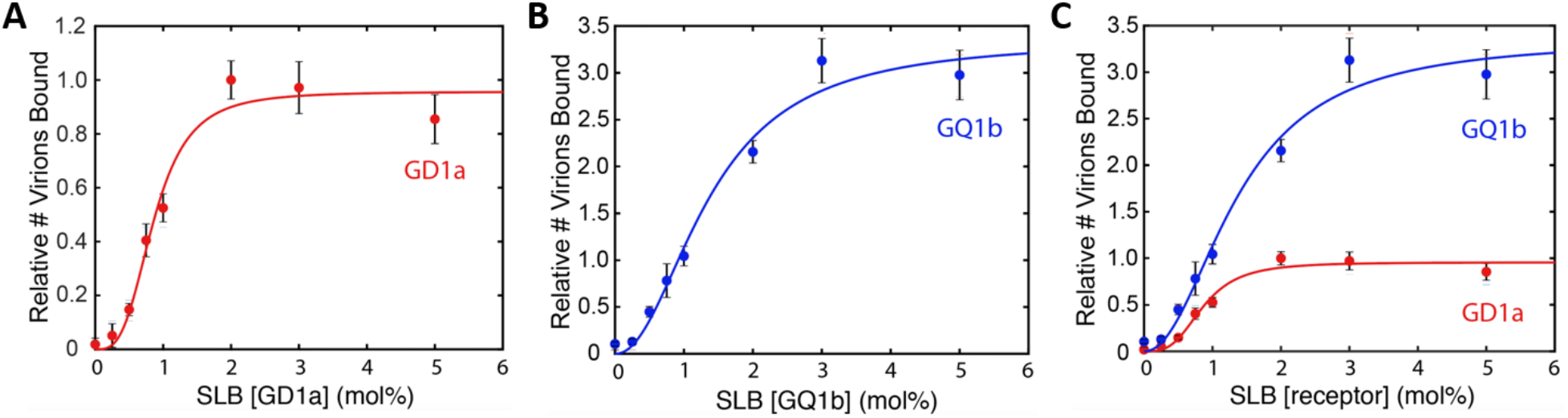
SeV exhibits cooperative binding to SLBs with either GD1a or GQ1b. Single virus binding measurements were performed to SLBs with varying concentrations of either A) GD1a or B) GQ1b. Solid lines show fits of the data to a Hill model binding curve (Eq. 1) to quantify the extent of cooperativity. The relative number of virions bound in both panels is calculated relative to 2% GD1a, and for ease of comparison are shown overlaid in Panel C. Error bars are ± standard error of ≥ 3 sample replicates, and ≥ 10 separate image locations within each sample. Best fit parameters are shown in Table 1.

where *RB*_*Max*_ is the maximum relative binding, *[receptor]* is the mole percent of receptor in the target SLB, *K* is equal to *[receptor]* when *RelBinding* = 0.5 x *RB*_*Max*_, and *C*_*F*_ is the cooperativity factor. Best fit values are shown in Table 1.

**Table 1.** Hill model fit values for GD1a and GQ1b cooperative binding data in Figure 3.

| Ganglioside Receptor | $RB_{max}$ | $K$ | $C_F$ |
| --- | --- | --- | --- |
| GD1a | $0.95 \pm 0.08$ | $0.86 \pm 0.07$ | $3.3 \pm 0.4$ |
| GQ1b | $3.6 \pm 0.80$ | $1.5 \pm 0.5$ | $2.1 \pm 0.4$ |

<sup>a</sup> All error estimates are determined by bootstrap re-sampling of individual samples and all images taken within each sample.

Interestingly, this best fit is quite similar to data collected for influenza virus A/H3N2/X-31 (IAV^X31^), which also can use GD1a as a receptor. In a similar binding experiment using SLBs with GD1a, best fit values for IAV^X31^ were *K* = 0.85, and *C*_*F*_ = 2.36 ± 0.85 (23), suggesting that both IAV^X31^ and SeV have evolved a similar cooperative binding response to sialic acid receptors, in spite of numerous structural and functional differences between the hemagglutinin of IAV^X31^ and the HN SeV. In protein enzymology, it has been noted that enzymes often have evolved a binding affinity which matches the physiological concentration of the ligand, presumably driven by selection to maximize their dynamic range (45). A similar driving force may be at play in the evolution of these two viruses.

We also studied the cooperative binding response of SeV using GQ1b as a receptor (Figure 3B). We observed a more pronounced curve relative to GD1a, with increased viral binding at all GQ1b concentrations > 0.25 mol%, consistent with the higher potency of GQ1b as a receptor compared to GD1a in bulk infectivity measurements (16). Best fit values to the Hill binding curve are reported in Table 1. The cooperativity factor for GQ1b was significantly lower than the cooperativity factor for GD1a (p-value = 0.03 by bootstrap resampling), whereas both *RB*_*Max*_ and *K* (half max concentration) were significantly larger for GQ1b than for GD1a (p-values = 0.006 and < 0.002, respectively). This suggests that fewer HN-GQ1b complexes may be needed for stable binding as compared to HN-GD1a complexes. This is likely due to the increased number of both internal and terminal sialic acid residues per ganglioside of GQ1b compared to GD1a (see Figure 2A).

### Cholesterol concentration in target SLB does not strongly influence viral binding

In studies of SeV and closely related paramyxovirus, fusion and infectivity are sensitive to cholesterol concentration in both the viral and target membranes, generally with less cholesterol resulting in less fusion (46–50). However, it is not clear whether this effect is due to modulation of binding, fusion, or both. In similar binding measurements to those performed here for SeV, it has been shown that cholesterol in the target SLBs can modulate binding of influenza virus (IAV^X31^) to GD1a (23), with ∼2 fold linear increase in binding avidity with increasing cholesterol concentrations from 0-30%. Using molecular dynamics simulations, it was suggested that this was due to the formation of receptor nanoclusters in the target SLB, which were stabilized by increasing concentrations of cholesterol. In the context of multivalent, cooperative binding, nanoclusters could function as binding “hotspots” to increase binding avidity.

Therefore, to examine whether cholesterol concentration in the target membrane would modulate SeV binding, we studied SeV binding to SLBs with either 1% or 2% GD1a and varying concentrations of cholesterol from 0-30%. The remaining lipid composition of the SLB was matched to what had been used for IAV^X31^. Interestingly, we observed that there was no strong dependence of SeV binding on SLB cholesterol concentration (see Figure 4). This suggests that sensitivity of SeV fusion and infection to cholesterol concentration is likely not due to viral binding with a similar mechanism IAV^X31^.

**Figure 4.**
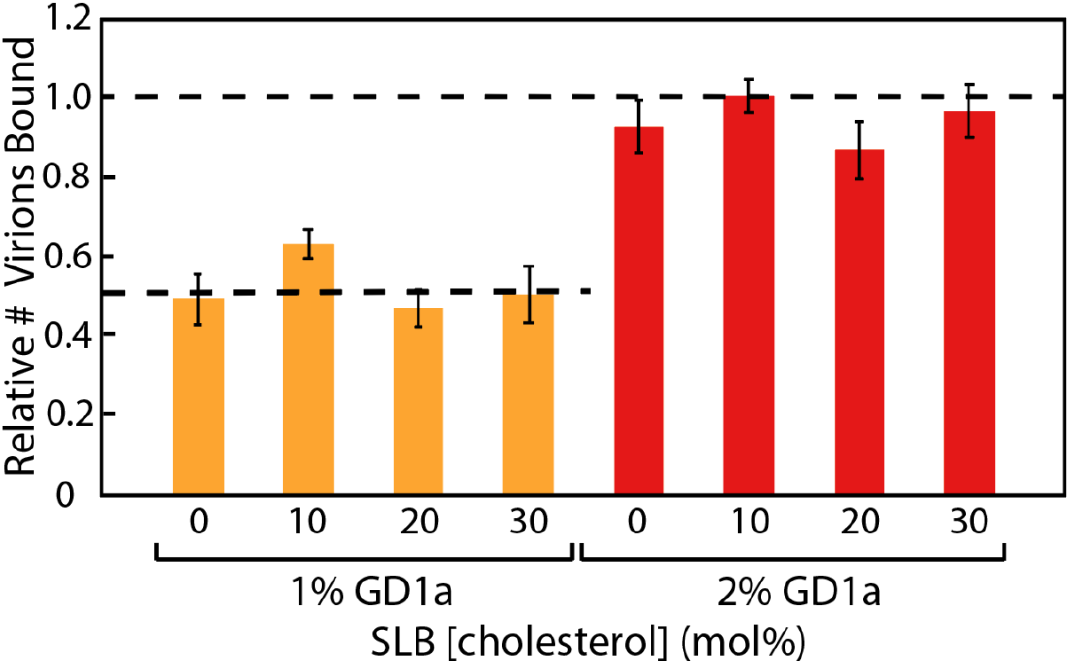
Sendai virus binding to SLBs with GD1a is insensitive to cholesterol concentration. Single virus binding measurements were performed to SLBs with varying concentrations of cholesterol and either 1% GD1a (yellow bars) or 2% GD1a (red bars). The SLBs also contained 20% DOPE, 0.05% Oregon Green-DHPE, and the remainder POPC (47.95-78.95%). The relative number of virions bound was calculated relative to 2% GD1a, 10% cholesterol. Within a set concentration of receptor, little difference in binding was observed across the range of cholesterol concentrations tested. By comparison, influenza A virus in ref (23) shows a ∼2 fold linear increase in binding to similar SLBs across the same range of cholesterol. Error bars are ± standard error of ≥ 3 sample replicates, and ≥ 10 separate image locations within each sample.

Surprisingly, this also suggests that SeV binding, unlike IAV^X31^, is insensitive to the formation of receptor nanoclusters in the target SLB, even though both viruses exhibit a similar cooperative binding dependence on GD1a concentration (see Figure 3 and discussion above). This indicates that while both viruses may have evolved a similar cooperative binding response to sialic acid-glycolipids, the binding mechanisms are distinct enough to produce different sensitivities to target cholesterol composition.

### Estimating τ_unbind_ for Sendai virus-SLB binding

To characterize SeV behavior and stability following binding, we performed time-lapse imaging of bound viral particles following solution rinse, quantifying net change in bound virions over time. We observed that bound viruses were quite stable on 2% GD1a membranes, decreasing only to ∼60-80% of their initial bound value over 60 minutes (Figure 6A). During this time, average SLB background intensities remained quite low, indicating that viral detachment, not membrane fusion, was the primary cause of the decrease. Detached viral particles could also be observed in the solution above the SLB.

**Figure 6.**
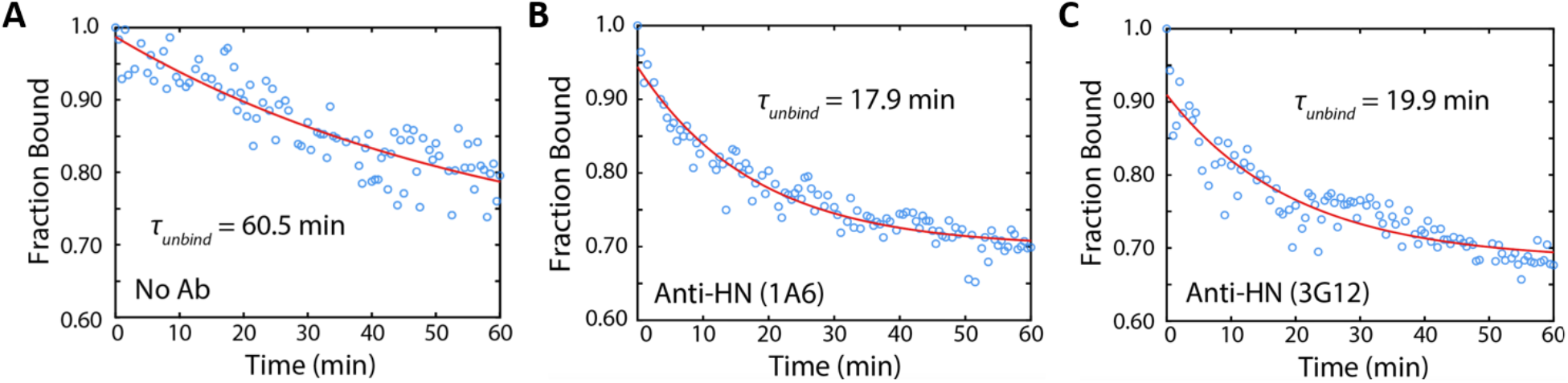
Antibody treatment following Sendai virus binding increases the rate of viral unbinding. Shown are example unbinding curves for SeV bound to SLBs with 2% GD1a. Following virus binding, antibody solution (or a no antibody control solution) was introduced into the flow cell, and viral unbinding from the SLB was observed over 1 hour. Shown are example data of the fraction of viruses bound over time in the presence of A) no antibody, B) anti-HN (1A6) at 20 µg/mL, and C) anti-HN (3G12) at 20 µg/mL. Red lines represent best fits to an exponential decay curve (Equation S4) to estimate the characteristic decay time (τ_unbind_).

To estimate the characteristic decay time of virus unbinding *τ*_*unbind*_, we fit the curves to an exponential decay (Equation S4), obtaining a *τ*_*unbind*_ of 79 ± 27 min. This indicates that once virus particles are bound to the underlying SLB, they remain quite stably attached for some time. Additionally, it is important to note that the SA receptor is in large excess on the SLB surface in these measurements (approximately 2 × 10^4^ per µm^2^ at 2% GD1a if we estimate 1 × 10^6^ lipids total per µm^2^) and that these measurements were made under non-optimal conditions (high Cl^-^ concentration, neutral pH, RT) for SeV neuraminidase activity (51). Therefore, the measured *τ*_*unbind*_ is unlikely to be simply due to the neuraminidase action of HN globally depleting the sialic acid density on the membrane over the course of the measurement.

### Virus mobility and evidence for “rolling” mechanism

We observed that many particles exhibited mobility by diffusion on the SLB following binding. To quantify the diffusion behavior, we performed single particle tracking of virions in time-lapse micrographs over 25 min and then estimated the diffusion coefficient (D) of each particle via linear fits of mean squared displacement (MSD) versus Δt curves. We observed that ∼45% of particles were immobile (D < 6 × 10^-5^ µm^2^/s, the lower limit of detectable diffusion). The majority of mobile particles exhibited low diffusion coefficients (D < 0.1 µm^2^/s), but a small percentage did exhibit higher values ranging up to 1.6 µm^2^/s (Figure 7). By comparison, 100 nm liposomes anchored by a single lipid in an SLB with a similar composition have D ≈ 1 µm^2^/s (52).

**Figure 7.**
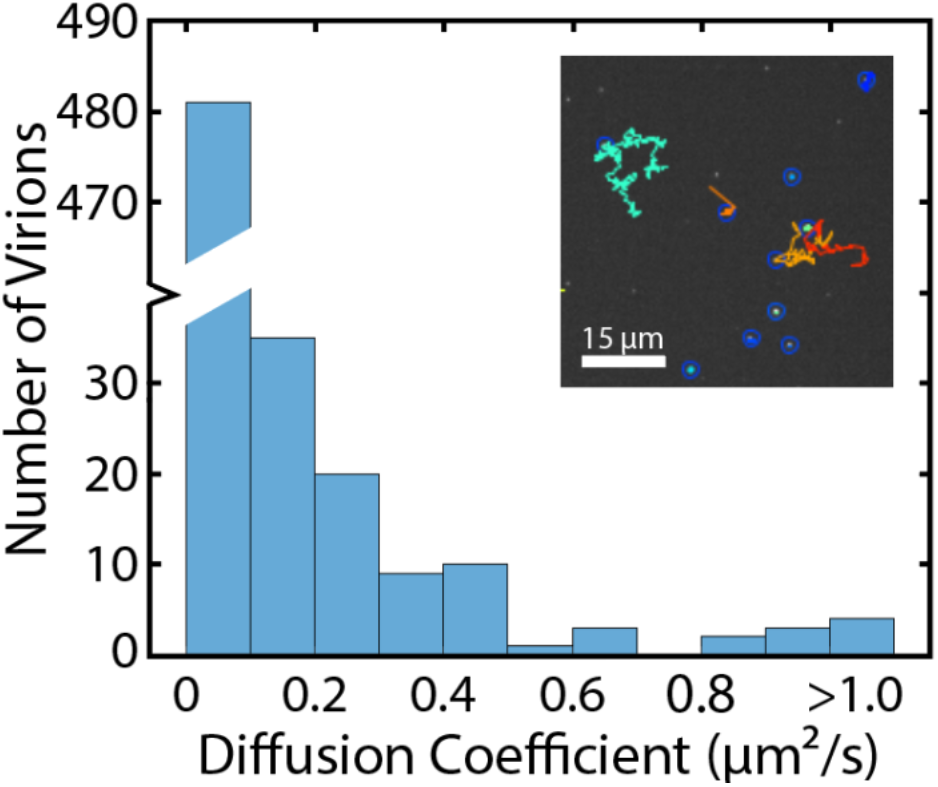
Distribution of diffusion coefficients for mobile virions on supported lipid bilayers. Diffusion coefficients were calculated from single particle tracking (SPT) traces of bound virions in time-lapse micrographs over 25 minutes. Inset shows sample SPT traces. 45% of particles were immobile (D < 6 × 10^-5^ µm^2^/s, the lower limit of detectable diffusion); these are not shown in the distribution. The vast majority (∼80%) of mobile virions exhibited slow diffusion D < 0.1 µm^2^/s. The SLB contained 2% GD1a. Total number of particles analyzed = 1035.

We anticipate two possible mechanisms for the observed diffusion of SeV on SLBs. First, SeV diffusion could be dominated by the diffusion of several lipid receptors to which the virus is stably bound. However, monomeric HN binding affinity for sialic acid-receptors is rather weak for closely related paramyxoviruses (∼0.1 mM *K*_*d*_ for HPIV-1, -2 and -3) and monomeric *k*_*off*_ values quite fast (∼0.2 s^-1^) as measured by surface plasmon resonance (44), suggesting that individual HN-receptor attachments are unlikely to be stably bound during our observation window. Instead, a more likely mechanism is a “rolling” mechanism, wherein numerous HN attachment proteins engage weakly with individual lipid receptors, and random binding and unbinding (and/or neuraminidase cleavage of the sialic acid) leads to Brownian diffusion on the SLB surface, as suggested for measurements of influenza virus (53, 54).

To differentiate between these mechanisms, we performed an antibody-mediated unbinding experiment (Figure 6B and C). Following viral binding and buffer rinse, anti-HN antibody (1A6) at 20 µg/mL was introduced into the flow cell and *τ*_*unbind*_ was measured as above. If viral diffusion were dominated by a “rolling” mechanism, we would expect a significant decrease in *τ*_*unbind*_, as the rolling particle became sequentially bound by a higher density of antibody until it could no longer sustain sufficient HN-receptor contacts to remain bound. If, on the other hand, HN-receptor complexes were stable and viral diffusion were dominated by lipid diffusion of receptors bound by HN, we would not expect a significant change in *τ*_*unbind*_, as the HN-receptor interface would be inaccessible to antibody binding; antibodies would predominantly bind to the distal viral surface. We observed that there was a significant decrease in *τ*_*unbind*_ following antibody treatment (Table 2), lending support to the “rolling” mechanism of viral diffusion.

**Table 2.** τ_unbind_ values^a^ for Sendai virus unbinding from SLBs in the presence or absence of anti-HN antibodies.

| Antibody Treatment | $\tau_{unbind}$ (min)<sup>b</sup> |
| --- | --- |
| No antibody | $79 \pm 27$ |
| Anti-HN (1A6) | $12.0 \pm 6.9^c$ |
| Anti-HN (3G12) | $25.6 \pm 7.6^c$ |

<sup>a</sup> The entire set of best fit values to the exponential decay curve is given in Table S1.
<sup>b</sup> Error estimates are the standard deviation of best fits to at least 3 sample replicates.
<sup>c</sup> Both the 1A6 and 3G12 $\tau_{unbind}$ values are significantly different than the No Antibody values ( $p < 0.015$ , 2-Sample Kolmogorov-Smirnov test).

What might be the mechanistic origins of the immobile virions? Several non-mutually exclusive explanations exist. First, immobile particles may be virions with a large enough number of HN-sialic acid engagements that “rolling” diffusion becomes difficult to observe at our experimental timescale (∼25 min). Tantalizingly, these particles may represent virions with large organized patches of HN on their surface, as has been reported in a cryo-EM study of HPIV3 (55), but for which no structural data as yet exists for SeV. Second, while all SLBs used in these measurements were observed to be defect-free by homogeneous fluorescence, we cannot rule out that some immobile particles may be bound to small, unobserved (sub-diffraction limit) defects. Third, immobile particles may be those with multiple activated F proteins, whose fusion peptides have been inserted into the SLB, but not yet collapsed into a post fusion conformation. However, as we observe little evidence of fusion, this would suggest that collapse of this unstable intermediate would be slow relative to the experiment timescale, which seems less likely.

## CONCLUSION

In this report, we have presented the development of a single virus binding assay for the prototypical paramyxovirus, Sendai virus, using glass-supported lipid bilayers as the target membrane for binding, and have discussed key design features that are critical to avoid unwanted artifacts. Using this binding assay, we investigated various biophysical questions relating to SeV binding. First, we found that the number and positioning of α2,3-linked sialic acids on receptors can strongly influence SeV binding. Second, we investigated the cooperative nature of SeV binding, observing a strikingly similar cooperative dependence on GD1a concentration with influenza A virus (IAV^X31^), suggesting a common evolutionary response toward receptor engagement despite many differences at the viral protein level. Third, we demonstrated that SeV exhibits little dependence on cholesterol concentration in the target SLB, suggesting that the sensitivity of SeV infection to cholesterol is likely due to a later step in the entry mechanism, and that unlike IAV^X31^, SeV binding is relatively insensitive to the presence of receptor nanoclusters. Finally, we characterized SeV behavior following binding, including the rate of detachment and viral mobility, and provided evidence for a “rolling mechanism” of mobility on the SLB surface. Together, these results provide mechanistic insight into the biophysics of SeV binding and may be useful to suggest constraints on host cell tropism and possible inhibitor design. We look forward to the extension of this assay to study other paramyxoviruses in the future.

## MATERIALS AND METHODS

### Materials

Dioleoylphosphatidylethanolamine (DOPE), palmitoyl oleoylphosphatidylcholine (POPC), and cholesterol (Chol), as well as gangliosides GD1a, GM3, GM1 and GQ1band cholesterol were purchased from Avanti Polar Lipids (Alabaster, AL). Oregon Green-1,2-dihexadecanoyl-sn-glycero-3-phosphoethanolamine (OG-DHPE) and Texas Red-1,2-dihexadecanoyl-sn-glycero-3-phosphoethanol-amine (TR-DHPE) was obtained from Thermo Fisher Scientific (Waltham, MA, USA). Polydimethylsiloxane (PDMS) Sylgard 184 elastomer base and curing agent were purchased from Ellsworth Adhesives (Germantown, WI, USA). Octadecyl rhodamine B chloride (R18) dye was purchased from Biotium, Inc. (Fremont, CA, USA). Sendai virus (purified Sendai Cantell Strain, batch 960216) was obtained from Charles River Laboratories (Wilmington, MA, USA) and handled according to a BSL-2 protocol at Williams College. Chloroform, methanol, and buffer salts were obtained from Fisher Scientific (Pittsburgh, PA) and Sigma-Aldrich (St. Louis, MO, USA). Formvar/carbon-coated square mesh grids (Cu, 200 Mesh, SB) were purchased from Electron Microscopy Sciences (Hatfield, PA, USA). Goat anti-Mouse IgG-Alexa 488 antibody was purchased from Abcam (Cambridge, United Kingdom).

### Anti-SeV Antibodies

The mouse anti-HN (1A6) IgG2a antibody and the mouse anti-HN (3G12) IgG1 antibody were ordered from Kerafast Inc. (Boston, MA), and produced in the laboratory of Prof. Benhur Lee (Mt. Sinai). Details of antibody production and characterization of 3G12 have not been published previously. Information on characterization of both monoclonal antibodies are included in the Supporting Information (Figure S9). The anti-F (11H12) monoclonal antibody was a gift from Prof. Benhur Lee. 1A6 and 11H12 have been characterized previously (42).

### Buffer Definitions

Reaction buffer (RB) = 10 mM NaH_2_PO_4_, 90 mM sodium citrate, 150 mM NaCl, pH 7.4. HEPES Buffer (HB) = 20 mM HEPES, 150 mM NaCl, pH 7.2.

### Sendai virus labeling and concentration estimation

Sendai virus (Cantell strain) was egg-grown and stringently purified by the manufacturer (Charles River Laboratories) using standard purification techniques (ultracentrifugation through sucrose gradient and tangential flow filtration). Viral particles were characterized by transmission electron microscopy (TEM), see details in SI. They were observed to be pleiomorphic, roughly spherical particles, with outer diameters ranging from 80 to 400 nm, mean 165 nm (Figure S10), consistent with more limited cryo-EM data in a prior report (58). To label virus, 0.5 μL (0.5X), 1 μL (1X), 2 μL (2X), or 4 μL (4X) of R18 (1.5 g/L in ethanol) OR 2 μL (0.5X), 4 μL (1X), 8 μL (2X), 12 μL (3X) or 16 μL (4X) of TR-DHPE (0.75 g/L in ethanol), was mixed with 240 μL of HB buffer. 15 μL of Sendai Virus (2 mg/mL) was combined with 60 μL of the appropriate dye buffer mixture, and this solution was incubated at room temperature for 2 hours. 1300 μL of HB was then added, and the solution was centrifuged for 50 min at 21K x g at 4°C to remove unincorporated dye. The supernatant was discarded and the pellet was resuspended in 100 μL HEPES buffer. Unless noted otherwise, the standard virus labeling was performed using 1X concentration of TR-DHPE. Labeled particle concentration was estimated either by viral protein content via BCA assay, or by spot counting on the microscope (see details in SI). Unless noted otherwise, values reported are calculated from BCA assay.

### Single virus binding and unbinding assay

Single virus binding assays were performed to SLBs formed by the vesicle fusion method inside microfluidic devices. SLB formation and microfluidic device construction have been described previously, and are outlined briefly in the Supporting Information. Unless otherwise noted, the standard SLB composition was 20 mol% DOPE, 10% Chol, 0.05% Oregon Green-DHPE, 0-5% ganglioside receptor (always listed), and the remaining amount (69.95-64.95%) POPC. Following SLB formation, 4 µL of labeled virus (typically at 0.1 nM concentration) was added to the inlet hole, and then drawn through to the outlet hole by pipette. Virus was incubated above the SLB for 15 minutes at RT (∼22°C). The flow cell was then rinsed with 1 mL RB by a syringe pump at 0.8 mL/min. Images of bound virions and SLB were then collected in 5-15 separate areas in the center region of the flow cell. Little variation in viral binding numbers along the length of the flow cell was observed, indicating that volume displacement during viral introduction was quite even. See Figure S2 for comparison with an alternate method of introduction. Viral spots in each field of view were quantified using previously published custom-built Matlab scripts (20, 21). Our version of these scripts, including parameters and options optimized for our Sendai virus data is available at https://github.com/rawlelab/SendaiBindingAnalysis. To measure viral unbinding, time-lapse fluorescence micrographs were taken every 30 seconds for 60 minutes following buffer rinse. For antibody-mediated unbinding measurements, 4 µL of 20 µg/mL antibody solution in RB buffer was introduced into the flow cell following buffer rinse, and then viral unbinding was observed as above.

### Antibody inhibition assay

For antibody inhibition measurements, labeled SeV was pre-mixed with antibody (Ab) solution; final concentrations of Ab as indicated. SeV-Ab mixture was incubated on ice for 30 min prior to introduction into the flow cell. Single virus binding measurements were then performed as above.

### Immunofluorescence measurement

Following SeV binding to SLB as above, 4 µL of anti-HN (1A6) at 20 µg/mL was introduced into the flow cell and incubated for 15 min. The flow cell was rinsed by syringe pump with 2 mL HB and then 4 µL of goat anti-mouse IgG-Alexa 488 at 2 µg/mL was introduced and incubated for 15 min. The flow cell was rinsed by syringe pump with 3 mL HB, and then sequential images of TR-labeled viral particles and Alexa 488-antibody were observed by fluorescence microscopy. Co-localization was quantified using previously published Matlab scripts (20, 21).

### Single Particle Tracking

For single particle tracking, time-lapse fluorescence micrographs of Sendai virions bound to SLBs were taken every 5 seconds for 25 minutes. Single particle tracking was performed using the TrackMate (56) plug-in of ImageJ. Image stacks were analyzed using a Laplacian of Gaussian (LoG) detector, which applies a LoG filter to the images with a sigma suited to the estimated blob size, 3.0 μm, and performs calculations in the Fourier space. In the median filtered image, the maxima were detected and those too close together were suppressed. Finally, a quadratic fitting scheme allowed for subpixel localization. The spots and tracks were then displayed using a HyperStack displayer, allowing for manual editing and filtering of spots by mean intensity (>420-445). Next, the virus particles, or spots, were tracked using the simple Linear Assignment Problem (LAP) tracker with a linking max distance of 8.0 μm, a gap-closing max distance of 8.0 μm, and a gap-closing max frame gap of 3.0 frames. Subsequently, the tracks were filtered by the number of spots per track (>100-200 spots per track) before manual editing. Manual editing was used to identify erroneous tracks (primarily those which the algorithm incorrectly assigned to jump back-and-forth between 2 adjacent immobile particles). The resulting MSD vs Δt curves were fitted to a linear model to extract the diffusion coefficients, following a published algorithm to determine the optimal number of MSD points to include in the fit (57). The Matlab code of our implementation of this algorithm is available online at https://github.com/rawlelab/SendaiBindingAnalysis. The lower limit of detectable diffusion (6.05 × 10^-5^ μm^2^/s) was determined by measuring the “diffusion coefficients” of virions bound directly to a glass surface with the single particle tracking method described.

### Fluorescence Microscopy

Fluorescence microscopy images were acquired with a Zeiss Axio Observer 3 microscope using a 63x oil immersion objective, NA=1.4 (Carl Zeiss Microscopy, LLC., White Plains, NY), with a Lumencor Spectra III, LED Light Engine as an excitation light source. Images were recorded with a Hamamatsu ORCA Flash 4.0 V2 Digital CMOS camera (Hamamatsu Photonics K.K., Hamamatsu City, Japan) using a 16-bit image setting and were captured with Micromanager software (Vale Lab, UCSF). Images and video micrographs were captured at 100 ms/frame with 2×2 binning. R18 and Texas Red images were obtained using the following filter cube: ex = 562/40 nm, bs = 593 nm, em = 641/75 nm, light engine typical intensity setting = 10/1000 (green LED). Alexa-488 and Oregon Green images were captured using the following excitation/emission filter cube: ex = 475/50 nm, bs = 506 nm, em = 540/50 nm, light engine typical intensity setting = 4/1000 (cyan LED).

## ACKNOWLEDGEMENTS

The authors thank Abraham Park (Williams College) and Elizabeth Webster (Sandia National Laboratories) for helpful manuscript feedback. RJR thanks Williams College for financial support. AL was supported by a Roche and Gomez Student Research Fellowship at Williams College. PAT was supported by a Canadian Institutes of Health Research postdoctoral fellowship. KDA acknowledges support from T32 AI007647-16 (Viral-Host Pathogenesis Training Grant at ISMMS) and NIAID F31-AI133943 (NIH Ruth L. Kirschstein Predoctoral Individual National Research Service Award). BL acknowledges support from NIH grant AI123449 and the Center for Therapeutic Antibody Development (CTAD) at the Icahn School of Medicine at Mount Sinai. The authors declare no competing interests.

## AUTHOR CONTRIBUTIONS

AL and OK designed experiments, collected and analyzed data, and helped write the manuscript. NS and DV collected data and contributed to single virus assay design. PFA collected bulk fusion data. PAT and KDA produced antibodies, and collected antibody characterization data. BL provided anti-SeV antibodies and antibody characterization data. RJR designed experiments, analyzed data, and wrote the manuscript.

