## Supplemental Information for "Single virus assay reveals membrane determinants and mechanistic features of Sendai virus binding"

### SUPPORTING MATERIALS AND METHODS

**PDMS Flow Cell Preparation**

PDMS flow cells were prepared using the "tape-based soft lithography" method as previously described (1). Briefly, flow cell molds were made by taping small strips (2.5 mm x ~13 mm x 70 µm) of Kapton polyimide tape (Ted Pella Inc., Redding, CA) to a glass microscope slide inside a petri dish. The PDMS mixture was prepared by weighing out a mixture of elastomer base and curing agent in a 10:1 ratio, degassed under vacuum for ~1 hour, and then poured to cover the entire petri dish. The mixture was then cured at 70°C for 2 hours. When solid, flow cells were cut out using a scalpel, and inlet/outlet holes created with a biopsy hole puncher (2.5 mm diameter, Harris Uni-core, Ted Pella Inc.). Flow cell volume was ~4 µL. PDMS flow cells were stored in a petri dish at room temperature with a small piece of scotch tape to protect from dust on the channel side. They were used for several weeks.

**Glass coverslip cleaning**

Glass coverslips (24 x 40 mm, No. 1.5 VWR International, Randor, PA) were cleaned in a ceramic holder by immersion in a 1:7 solution of 7x detergent (MP Biomedicals, Burlingame, CA) and deionized water, with heating until the solution became clear. The coverslips were rinsed under running DI water for >2 hours, given a final rinse with Millipore water, and then baked in a kiln at 400°C for 4 hours. They were then stored at room temperature and used for several weeks.

**Microfluidic device assembly and supported lipid bilayer formation**

Supported lipid bilayers were prepared using the vesicle fusion method, as previously described (2). Briefly, a lipid mixture at the desired ratio of POPC, DOPE, cholesterol, OG-DHPE, and ganglioside receptor in 4:1 chloroform:methanol was prepared in a glass test tube, with 2.8 x $10^{-7}$ total moles lipid. Solvent was evaporated under $N_2$(g) to a lipid film, which was dried under house vacuum for at least several hours to remove any residual solvent. The lipid film was resuspended by vortexing in 250 µL of Millipore water, and extruded at least 20x using a mini-extruder (Avanti Polar Lipids, Alabaster, AL) with membrane pore size 50 or 100 nm to produce large unilamellar vesicles. Vesicles were stored at 4°C, and used within several days. Unless otherwise noted, the standard vesicle (and SLB) composition was 20 mol% DOPE, 10% cholesterol, 0.05% Oregon Green-DHPE, 0-5% ganglioside receptor (always listed), and the remaining amount (69.95-64.95%) POPC.

Immediately before SLB formation, a microfluidic device was assembled by plasma bonding. A PDMS flow cell (channel side facing up) and a clean glass coverslip were exposed to air plasma for 60 s (Harrick Plasma Cleaner PDC-3xG, Harrick Plasma, Ithaca, NY). After plasma exposure, the flow cell was immediately placed channel side down onto the glass coverslip for bonding. A vesicle suspension was mixed 1:1 with RB, and 5 µL was immediately added to each channel via the inlet holes. The vesicle suspension was incubated inside the channel for 30 minutes to allow SLB formation. Meanwhile, a buffer well cut from the end of a 1 mL plastic pipette tip was then affixed to the inlet hole of each channel with five-minute epoxy (Devcon, Danvers, MA). Channels were then rinsed with 1.5 mL of Millipore water and then with 1.5 mL RB using a Fusion 200 syringe pump (Chemyx Inc., Stafford, TX, USA), flow rate 800 µL/min. After rinsing, bilayers were imaged using the fluorescent microscope. Homogeneous fluorescence with little or no debris and defects was used as quality control to verify SLB integrity. SLBs were observed to be fluid by fluorescence recovery after photobleaching.

**Virus concentration determination**

Viral particle concentration was estimated via two different methods, both of which gave similar estimates.

The first method was based on an estimated particle count of $1.3 \times 10^9$ viral particles per µg of protein, previously reported in the literature for Sendai virus (3) and estimated from data of the chemical composition of Sendai virus. We measured the viral protein concentration in our samples following fluorescent labeling using a BCA assay with comparison to a BSA standard. A typical protein concentration used in our experiments was 50 µg/mL, which then corresponds to 0.11 nM viral particle concentration.

The second method, nicknamed the “splat assay”, used direct visualization of fluorescently labeled viral particles on a glass coverslip by fluorescence microscopy. Viral particles were diluted to an appropriate concentration to easily quantify individual spots and were pre-mixed with liposomes at a known nominal concentration. Liposomes contained 0.5% Oregon Green-DHPE for visualization in a separate fluorescent channel, but did not contain any viral receptor, preventing any virus-liposome binding or fusion. This virus-liposome mixture was then added to an empty flow cell, and was incubated for 30 minutes. Nearly all particles adhered to the coverslip surface during this incubation, as observed by the absence of floating particles in the solution above. Fluorescence micrographs were then acquired of the viral particles and the liposomes, and the number of each was quantified. The ratio of bound viral particles to bound liposomes within a field of view could then be used to estimate the viral particle concentration as follows:

$$ConcVirus = \frac{NumBoundVirus}{NumBoundLiposomes} \times ConcLiposomes$$

**Equation S1**

The known nominal concentration of liposomes (*ConcLiposomes*) was calculated from the lipid concentration used to prepare the liposomes (*ConcLipids)*, the average cross-sectional area of a lipid ($SA_{lipid}$), and the average surface area of a single liposome (calculated from the average diameter of the liposomes, $d_{liposome}$) multiplied by 2 to account for both leaflets.

$$ConcLiposomes = \frac{ConcLipids \times SA_{lipid}}{2 \times 4\pi \left(\frac{d_{liposome}}{2}\right)^2}$$

**Equation S2**

*ConcLipids* was calculated from the lipid concentration reported by the manufacturer, accounting for any dilutions during liposome preparation, and assuming no lipid loss during the preparation procedure. $d_{liposome}$ of a liposome was estimated to be 100 nm, the nominal extrusion pore size. The average cross-sectional area of a lipid, $SA_{lipid}$, was set at 60 Å$^2$ (4).

Using this method, we estimated viral particle concentration following labeling at 0.09 nM, remarkably close to the estimation by the first method. We note that this second method assumes that all fluorescently labeled particles are in fact viral particles. Our IFA results indicate that only ~30% of particles are IFA positive for surface glycoproteins. But we cannot rule out that the remaining particles may still contain viral protein, but simply lack a higher density of HN and F on their surface. In any case, the viral particle concentration estimations by both methods are in relatively close agreement.

**<u>Bulk fusion (lipid mixing) measurements</u>**

Bulk fusion measurements were based on those described in Hoesktra et al. (5). Lipid mixing was detected by relief of self-quenching of fluorescence upon fusion between labeled virus and unlabeled vesicles. Fluorescence of the reaction mixture was measured using a Perkin Elmer Fluorescence Spectrometer LS 55 with temperature control (Ex = 550 ± 10 nm, Em = 585 ± 20 nm). 200 µL of vesicles (3.17 nM total lipids) were added to 1800 µL RB buffer in a quartz cuvette and allowed to incubate for at least an hour. Then, 45 µL of labeled virus (1.2 µg of viral protein) was added to the cuvette while stirring and fluorescence was recorded for 1 hour. At the end of the reaction, 100 µL of 10% Triton-X detergent in RB buffer was added to the reaction mixture to completely solubilize the viral membrane so that maximum dequenching could be observed. Lipid mixing was quantified as relative percent fluorescence intensity (*Rel%F*):

$$Rel\,\%\,F = \frac{F_t - F_{t=0}}{F_{max} - F_{t=0}}$$

**Equation S3**

where $F_t$ is the observed fluorescence intensity at time t, $F_{max}$ is fluorescence intensity upon Triton-X treatment accounting for dilution, and $F_{t=0}$ is the initial fluorescence at t = 0 following virus addition.

Vesicle composition for all bulk lipid mixing measurements was 68% POPC, 20% DOPE, 10% Chol, and 2% GD1a. Virus was labeled with R18 at 1X concentration. Heat-inactivated virus was prepared by incubating virus at 70°C for 20 minutes immediately prior to lipid mixing measurement.

**<u>Viral unbinding analysis</u>**

Viral unbinding data (e.g. Figure 6) were fit to an exponential decay curve:

$$FracBound = A\,e^{-t/\tau_{unbind}} + C$$

**Equation S4**

where *FracBound* is the observed fraction of bound virus at time t, *A* is the scaling constant, $\tau_{unbind}$ is the characteristic decay time of unbinding, and *C* is the offset. Inclusion of an offset assumes that there is some population of virions which are permanently bound, or at least unbind with a very long decay time. This appears consistent with our data, which shows a visible plateau occurring in the antibody decay curves (e.g. Figure 6B and C in the main text).

Negative Stain Electron Microscopy

Sendai viral samples were prepared for negative-stain electron microscopy using the side blotting method (6, 7). Briefly, 5 µL aliquots of undiluted SeV were adsorbed onto formvar/carbon-coated square mesh grids by touching the coated side of each grid to the surface of the viral sample. The grids used for adsorption were either dry or pre-coated with a layer of ultrapure water. Grids were then allowed to stand for 1 minute and the excess sample was blotted onto filter paper. The samples were then stained twice for 30 seconds each with 50 µL of a 1% uranyl acetate solution and the excess blotted onto filter paper. The negative-stained grids were allowed to air-dry for 1 hr and then imaged using a JEOL JEM-1400 Plus transmission electron microscope operating at an acceleration voltage of 120 kV. Images were captured with a Gatan Orius SC1000A charge-coupled device (CCD).

**Antibody production and purification**

Hybridoma cultures for 1A6, 3G12, and 11H12 were originally produced from the fusion of primary mouse splenocytes, following live intraperitoneal SeV Cantell strain infection, with mouse myeloma cells.

The hybridoma cultures were initially expanded in Clonacell-HY Medium E but gradually switched to serum-free hybridoma medium until a final volume of 50 to 75 mL was achieved. When cells appeared no longer viable (~10 days after the final expansion step), the cultures were harvested by low-speed centrifugation (30 min, 5,500 x *g*), and the supernatants were passed through 0.22-µm-pore size sterile filtration units (Millipore).

Antibodies were purified using gravity flow over a bed of protein G Sepharose (GE Life Sciences). The column was washed with three column volumes (CVs) of phosphate buffered saline (PBS) before application of the supernatant. Flowthrough was run over the column an additional two times before washing with three CVs of PBS. Elution was performed into 2 M Tris·HCl, pH 10, by addition of 0.1 M glycine at pH 2.5 to the column at a ratio of 1:9. Eluate was stored at 4 °C until buffer exchange. Monoclonal antibody was buffer-exchanged against PBS using an Amicon Ultra-15 filter with 30-kDa cutoff (Mirus). Protein concentration was determined by Nanodrop absorbance at 280 nm.

**Antibody Characterization**

*Neutralization Assay:* 24h prior to infection, HeLa cells (cultured in DMEM + 10% FBS) were seeded in 96-well plates at a density of 2 x $10^5$ cells per mL, 100 µL cells per well. Antibody and virus (SeV, HPIV1, or HPIV3 (*human respirovirus 1* and *3*) encoding an eGFP fluorescent reporter gene (8) were combined in plain DMEM and incubated for 1h at room temperature, then culture media was removed from cells and replaced with antibody:virus inoculum. All virus inocula were carried out at a multiplicity of infection (moi) of 0.5, and all antibody:virus combinations were tested in technical triplicate. 24h after infection, cells were fixed with 2% paraformaldehyde in PBS, and infection was quantified using a Celigo plate imager to count infected cells by eGFP positivity. Percent virus inhibition for each antibody was calculated relative to HeLa cells infected in the absence of antibody.

*Hemagglutination Inhibition Assay:* SeV was first quantified by hemagglutination assay against guinea pig red blood cells (RBCs; Lampire Biologicals): serial two-fold dilutions of virus stock were combined with RBCs to a final concentration of 0.25% RBCs in HA buffer (Fisher Scientific) in a V-bottom 96-well plate, and incubated at 4°C for approx 5h. Hemagglutination was observed by the absence of an RBC "button" in the bottom of the well, and HU (hemagglutinating units) were defined by the reciprocal of the last dilution at which RBCs do not form a button (*i.e.* they are retained in suspension by interactions between virions and RBCs). To test hemagglutination inhibition activity (HAI), antibody was prepared in serial two-fold dilutions with 4 HU of SeV, and combined with RBCs to a final concentration of 0.25% RBCs in HA buffer, in a 96-well V-bottom plate. Preparation was incubated for 5h at 4°C, and then observed: HAI was defined by the lowest concentration of antibody at which RBCs still formed a "button" in the presence of virus.

*Flow cytometry:* BsrT7 cells were selected for stable integration of pCMV-Tet3G (Clontech) and a linearized resistance marker with puromycin (7 µg/mL) and single-cell cloned. The best-expressing clone was then further selected for stable and inducible expression of codon-optimized SeV-F in pTRE3G with hygromycin (100 µg/mL), and maintained in both selective antibiotics. SeV-F expression was induced with doxycycline at 1000 µg/mL for 24h. Parental cells with or without SeV-eGFP infection at an moi of 2, were used as controls.  24h post-induction or post-infection, cells were dissociated with PBS+20mM EDTA, pelleted, and fixed in 2% paraformaldehyde for 10 min. Cells were washed twice with FACS buffer (PBS + 2% FBS + 2mM EDTA), then incubated with indicated primary antibody at 1:20 in FACS buffer for 1hr. Cells were washed three times with FACS buffer, then incubated with secondary antibody (goat anti-mouse Alexafluor-647, Invitrogen) for 1h at 1:4000 in FACS buffer. Cells were washed three times, then analyzed for eGFP fluorescence (infection) and far-red fluorescence (antibody reactivity) using a Guava easyCite flow cytometer.

### SUPPORTING DATA

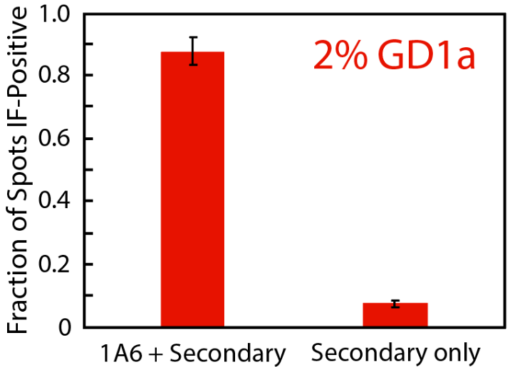


***Figure S1. Immunofluorescence imaging control experiment.*** *Plotted are the fraction of spots bound to SLBs 2% GD1a which show positive immunofluorescence (IF) labeling by anti-HN 1A6 antibody, followed by Alexa-488-labeled secondary antibody (1A6 + Secondary) or mock treated with HB buffer, followed by secondary antibody (Secondary only). Colocalization between Alexa-488 and the TR label in fluorescence micrographs was used to determine if a particle was IF-positive. Particles treated with only secondary antibody showed little colocalization, indicating little non-specific binding of the secondary. Error bars are ± standard error of 1-3 sample replicates with ≥ 10 separate image locations within each sample.*

#### Assay Development and Optimization

As discussed in the main text, we found that three critical aspects of our single virus assay design were 1) the method of introduction of the virus into the microfluidic chamber, 2) the extent of labeling and identity of the fluorescent dye in the virus, and 3) the minimization of receptor-triggered membrane fusion. Each of these is discussed below, together with supporting data.

1) ***Method of introduction***

   Where reported, previous single particle assays typically introduce viral particles either at low concentration under constant flow conditions, or by introduction at high concentration via pipette, followed by multiple rounds of mixing in the flow cell to ensure adequate mixing. For SeV, we found that the preferred method of introduction was a single, complete exchange of the flow cell volume with the viral suspension followed by incubation under stationary conditions. Mixing of smaller volumes within the flow cell led to much higher rates of non-specific binding and more sample-to-sample variability (Figure S2), presumably due to shear flow-mediated surface interactions during mixing.

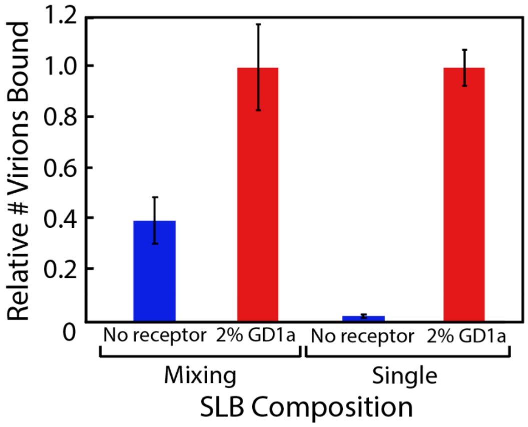


***Figure S2. Comparison of SeV binding data on method of introduction of virus into the flow cell.*** *Shown is the relative number of bound virions to SLBs either with 2% GD1a or without receptor, where the labeled virus was introduced into the flow cell either by a single, complete exchange of the flow cell volume by pipette or by multiple rounds of mixing of the viral suspension with the volume in the flow cell by pipette. The mixing method led to more variability (larger error bars) and to increased amounts of non-specific binding to SLBs without receptor than the single exchange method, presumably due to shear flow-mediated surface interactions during mixing. Values are shown relative to 2% GD1a for each method. Error bars are ± standard error of ≥ 3 sample replicates, and ≥ 10 separate image locations within each sample.*

2) ***Fluorescent labeling***

Fluorescence labeling of lipid membranes can, under some circumstances, lead to unwanted perturbations (9), and it has been demonstrated that influenza virus fusion kinetics can be altered by the lipophilic dye used to label the viral envelope (10). We investigated the effect of fluorescent labeling on SeV binding using 2 common dyes used to label viral envelopes – R18 and TR-DHPE (Figures S3-S4). We found that while SeV was quite sensitive to the concentration of R18 added, it was relatively insensitive to the concentration of TR-DHPE added. At low concentrations, both dyes exhibited roughly similar binding and sensitivity to antibody inhibition and so could be used interchangeably, but at higher concentrations, R18-labeled virions became de-stabilized and fused non-specifically with the supported bilayer. This sensitivity to R18 is of particular note because many reports studying SeV fusion to liposomes or erythrocyte ghosts in bulk have utilized R18 labeling (see for example refs (11–15)). Depending on the extent of labeling, non-specific fusion artifacts may have been occurring.

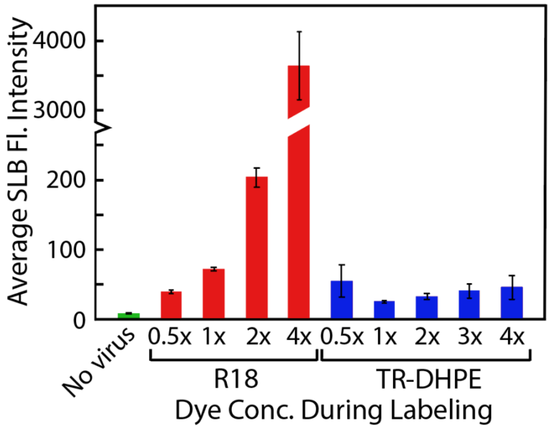


***Figure S3. Sendai virus is more easily de-stabilized by R18 concentration during labeling than Texas Red-DHPE.*** *Single virus binding assays were conducted at RT with SeV labeled with different self-quenching concentrations of R18 or TR-DHPE, and the average background SLB fluorescence intensity was quantified following virus binding and buffer rinse. If widespread lipid mixing occurred, the SLB intensity would increase as dye from the virions was transferred to the SLB and dispersed by diffusion. Average SLB intensity was calculated by averaging all pixels in a micrograph after excluding bound virions. SeV labeled with R18 was quite sensitive to the dye concentration added, with higher concentrations leading to de-stabilization and non-specific fusion. Conversely, SeV labeled with TR-DHPE exhibited low background intensities across all concentrations tested. Note that the Y-axis is broken for ease of visualization. Error bars are ± standard error ≥ 10 separate image locations within each sample.*

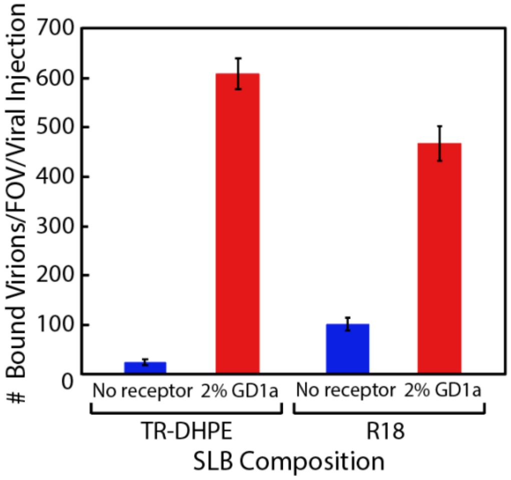


***Figure S4. Comparison of observed numbers of bound virions for SeV labeled with either TR-DHPE or R18 at a sub-destabilizing concentration (1X).*** *Virus was labeled at 0.5x and 1X concentration for R18 and TR-DHPE, respectively, and then a side-by-side single virus binding assay was performed to SLBs with either 2% GD1a or no receptor. The amount of virus added was standardized by viral protein concentration at 50 µg/mL. Shown is the number of bound virions per field-of-view (FOV, 211 x 211 µm) per standardized viral injection into the flow cell. Roughly similar binding results were observed for both dyes, although R18 exhibited slightly less binding to 2% GD1a SLBs and slightly more non-specific binding to SLBs without receptor. Error bars are ± standard error of ≥ 3 sample replicates, and ≥ 10 separate image locations within each sample.*

3) ***Minimizing membrane fusion.*** Many single virus assays utilize viruses for which fusion is triggered by pH or other externally added factors which can be manipulated separately from binding. However, for paramyxoviruses like SeV, receptor binding by HN can trigger the viral F protein to initiate membrane fusion (16). This can confound accurate quantification of viral binding, and therefore was something we wished to minimize in our assay. Previous research has demonstrated that Sendai fusion is temperature sensitive, with robust fusion occurring in bulk experiments only above 30-40°C (17, 18). In our own fusion assays monitoring lipid mixing between fluorescently labeled Sendai virions and target liposomes in bulk, we observed that lipid mixing at room temperature (~22°C) was reduced to the level of heat-inactivated virus, substantially lower than lipid mixing at 37°C (Figure S5). Therefore, to study virus binding in the absence of membrane fusion, we performed our single virus fusion measurements at room temperature.

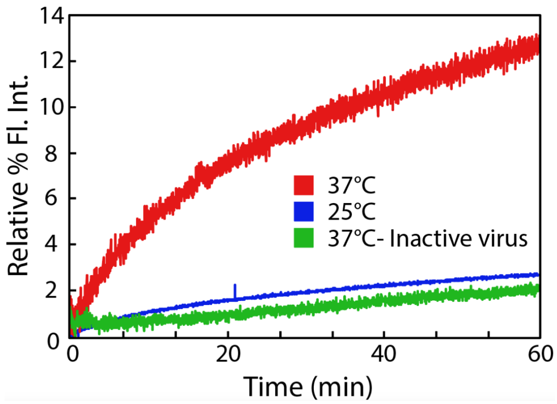


***Figure S5. Bulk lipid mixing measurements indicate that SeV is largely fusion inactive at 25°C.*** *45 µL of R18-labeled virus (1X R18 concentration, 1.2 µg of viral protein) was mixed with unlabeled vesicles with 2% GD1a (0.3 nM total lipid concentration) in a quartz cuvette and monitored by fluorescence spectroscopy for 1 hour at the temperature indicated (25°C or 37°C). Lipid mixing was detected as fluorescence de-quenching upon fusion between labeled virus and unlabeled vesicles. After 1 hour, 100 µL of 10% Triton-detergent was added to completely solubilize the vital membrane and observe maximum de-quenching. Relative % fluorescence intensity was calculated using this maximum value, as shown in Equation S3. Inactive virus was prepared by incubating virus at 70°C for 20 minutes immediately prior to the bulk lipid mixing measurement. Whereas lipid mixing at 37°C was robust for active virus, lipid mixing at 25°C was reduced to the level of inactive virus.*

To verify that this approach was successful in our single virus binding assay, we monitored lipid mixing as a marker for membrane fusion by incorporating a self-quenched concentration of lipophilic dye into the viral envelope. If lipid mixing with the target SLB occurred while imaging individual viruses, dye transfer to the SLB could be easily observed by a transient increase in the fluorescence intensity of the viral particle, followed by radial expansion of dye in the SLB due to diffusion (see Movie S1 and Figure S6). Additionally, widespread lipid mixing of many viral particles over time could be easily observed by an increase in the average background intensity of the SLB.

Using this approach, we rarely (<<1%) observed lipid mixing events for individual viral particles in the 15 minutes following binding, a typical timescale of our measurements. Over longer times, average background intensities also remained very low, unless the virus had been destabilized by too high concentration of dye (see above). Together, this data suggests that our assay design was successful at isolating virus binding from fusion. We also note that while the temperature during the measurement is an obvious factor in this observation, another intriguing factor for future study is the role of the SLB itself. In other membrane fusion systems, strong coupling between the lipid bilayer and the underlying glass support has been shown to inhibit membrane fusion (19, 20).

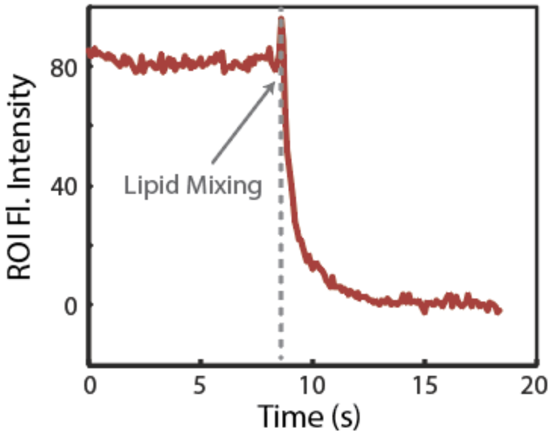


***Figure S6*. *Example lipid mixing trace of a Texas Red-labeled Sendai virion fusing with an SLB.*** *This trace corresponds to the virion which fuses in Movie S1. Plotted is the fluorescence intensity (relative units) within a region of interest (ROI) around the virion over time. Lipid mixing is observed as a small de-quenching spike, followed by rapid decay as the Texas Red-DHPE lipids diffuse into the SLB and out of the ROI. As discussed in the main text, these lipid mixing events were extremely rare (<<1%) in our assay, which was designed to study viral binding in the absence of fusion.*

Link to movie provided in Supplementary Material online

***Movie S1. Example lipid mixing event of a Texas Red-labeled Sendai virion fusing with an SLB.*** *In the movie, several virions bound to an SLB are observed. Some are mobile, some are not. One virion in the center of the field of view is observed to undergo lipid mixing in an "explosion", transferring its membrane dye to the SLB, followed by outward radial diffusion of the dye in the SLB. As discussed in the main text, such lipid mixing events were extremely rare (<<1%) in our assay, which was designed to study viral binding in the absence of fusion. For ease of viewing, the movie has been sped up 3.3X from the original data collection (collected at 100 ms per frame). The dimensions of the field of view are 37 x 38 μm.*

**<u>Immunofluorescence Imaging Results: SLB binding efficiently selects active virus particles</u>**

As mentioned briefly in the main text, viral particles were characterized following dye labeling by immunofluorescence (IF). Particles were bound non-specifically to a clean glass coverslip, and then labeled with anti-HN antibody (1A6) followed by an Alexa-488 secondary antibody. Although viral particles had been stringently purified during preparation using conventional methods (see Materials and Methods in the main text), we observed that only ~30% of particles were IF-positive (Figure 1D in the main text). This percentage of IF-positive particles is consistent with other single virus reports where such measurements have been reported (21). The other ~70% of non-IF positive particles likely represent extracellular vesicles or other cellular particles which are co-purified with the virions, are also labeled during dye incubation, and which may therefore be a confounding factor in single virus experiments. It is also possible that these particles may be viral intermediates, and include non-HN viral proteins.

To determine whether SLB attachment would selectively engage only the IF-positive particles, we also performed immunofluorescence labeling of viral particles following SLB attachment (Figure 1D in the main text). We observed that nearly all (> 85%) particles bound to SLBs with receptor (either GD1a or GQ1b) were IF-positive, and this did not depend on choice of labeling dye. However, particles bound to SLBs without receptors had a similar percentage of IF-positivity as particles bound non-specifically to glass. Together, these results suggest that binding to SLBs with receptor efficiently selects IF-positive viral particles, whereas the comparably lower binding to SLBs without receptor is largely dominated by non-specific interactions of any particles in the prep (viral or not).

Other reports have detailed the difficulties in separating extracellular vesicles or quasi-virus intermediates from virions (22–24), and our observation here serves as a potential warning for other researchers performing single virus measurements. Careful confirmation of selective virus binding using IF-labeling or other techniques is essential to validate that the results of the measurement are due to viral particles as opposed to other particles in the prep which can be difficult to separate.

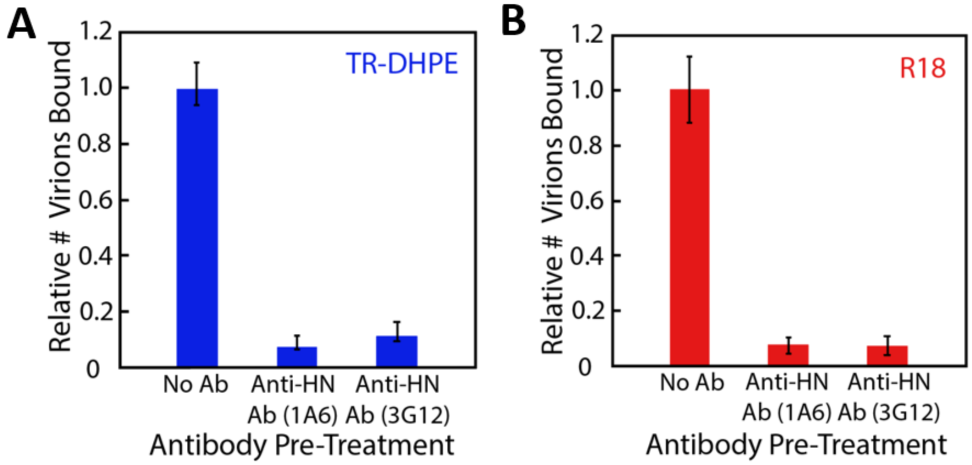


***Figure S7. Antibody inhibition of binding is similar regardless whether SeV is labeled with TR-DHPE (A) or R18 (B).*** *Single virus binding experiments to SLBs with 2% GD1a were performed in which virus was pre-treated with 25 µg/mL of monoclonal antibody (1A6 or 3G12) or a mock control solution (HB buffer) prior to injection into the flow cell. Virus was labeled at a sub-destabilizing concentration (1X TR-DHPE and 0.5X R18) for both dyes. Values are shown relative to the no antibody condition for each panel. Error bars are ± standard error of ≥ 3 sample replicates, and ≥ 10 separate image locations within each sample.*

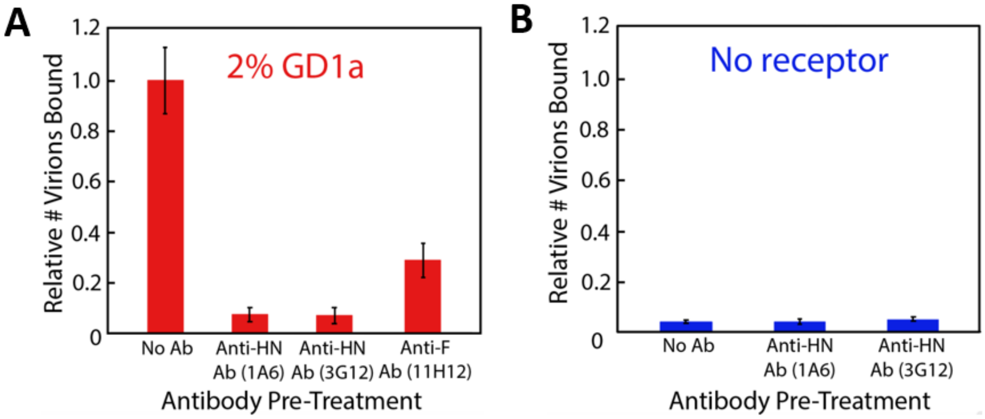


***Figure S8. Inhibition of SeV binding to SLBs by different monoclonal antibodies.*** *Single virus binding experiments were performed in which virus was pre-treated with 25 µg/mL of monoclonal antibody (1A6, 3G12, or 11H12) or a mock control solution (HB buffer) prior to injection into the flow cell. Shown are the relative numbers of bound virions to SLBs with 2% GD1a (A) or no receptor (B). For all antibodies tested, inhibition of viral binding to 2% GD1a SLBs was observed. Little difference was observed in binding to SLBs without receptor. All values are shown relative to the 2% GD1a, no antibody condition. Error bars are ± standard error of ≥ 3 sample replicates, and ≥ 10 separate image locations within each sample.*

***Table S1. Exponential decay fit values for Sendai virus unbinding from SLBs in the presence or absence of anti-HN antibodies using Equation S4***

| **Antibody Treatment** | $\tau_{unbind}$ **(min)** | **A** | **Offset** |
|---|---|---|---|
| **No antibody** | 79 ± 27 | 0.51 ± 0.19 | 0.48 ± 0.19 |
| **Anti-HN (1A6)** | 12.0 ± 6.9 | 0.18 ± 0.07 | 0.78 ± 0.07 |
| **Anti-HN (3G12)** | 25.6 ± 7.6 | 0.20 ± 0.07 | 0.74 ± 0.12 |

***Note 1:*** *Error estimates are the standard deviation of best fits to at least 3 sample replicates.*

### Monoclonal Antibody Characterization Data (1A6, 3G12, 11H12)

We first tested all antibodies for neutralization capacity against SeV, and against closely-related viruses HPIV1 and HPIV3 (human respiroviruses 1 and 3; Fig. **S9A-C**). No antibodies cross-reacted with HPIV1 or 3, and only 1A6 and 3G12 demonstrated neutralization activity ($IC_{50}$ 1.0 µg/mL and 1.7 µg/mL, respectively) against SeV. We also assayed all three antibodies for hemagglutination inhibition (HAI) against SeV to detect which antibodies interact directly with the receptor binding pocket of the SeV HN glycoprotein (Fig. **S9D**). 1A6 demonstrated strong HAI at 0.4 µg/mL, 3G12 showed weak HAI at 3.3 µg/mL, and 11H12 showed no HAI. Thus, 1A6 and 3G12 definitively bind SeV-HN. To identify the target of 11H12, we tested the antibodies by flow cytometry analysis of SeV-infected, non-permeabilized BsrT7 cells (Fig. **S9E-F**, green), then determined that it specifically bound SeV F (fusion glycoprotein) when we then assayed its binding of BsrT7 cells stably and inducibly expressing SeV F (Fig. **S9F**, orange). Finally, when we attempted to assay all three antibodies for western blot activity, we could not detect SeV glycoproteins by western blot (not shown) and determined that all three antibodies bound conformational epitopes of SeV.

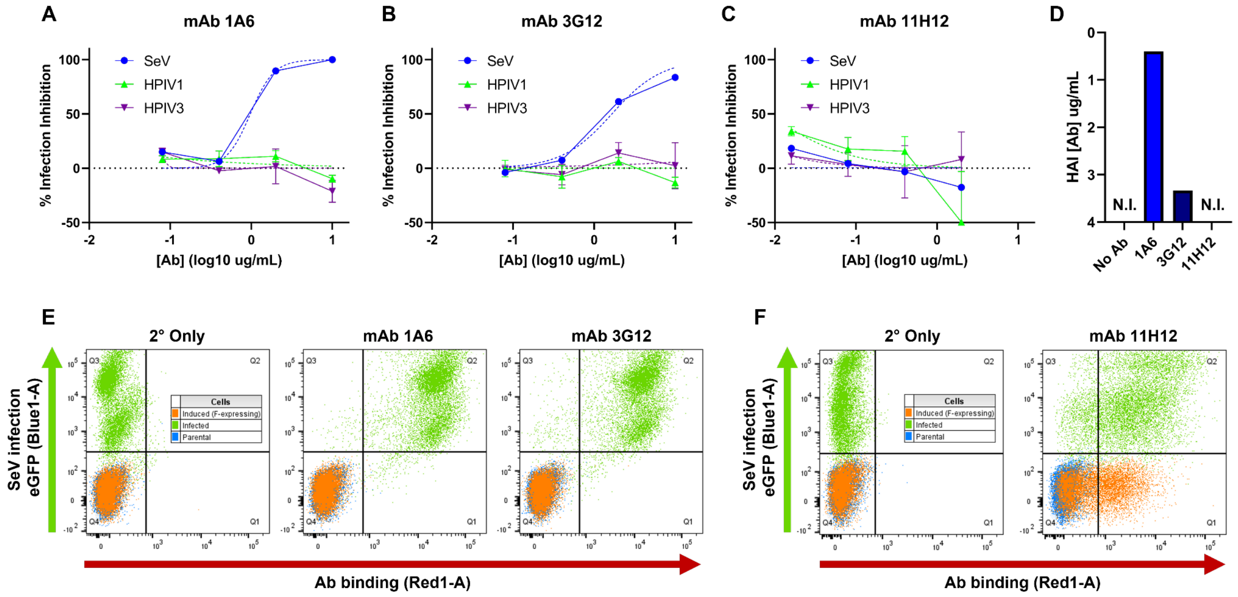

***Figure S9. SeV mAb characterization.*** *Antibodies 1A6 **(A)**, 3G12 **(B)**, and 11H12 **(C)** were serially diluted and tested for neutralization activity (% infection inhibition) against SeV (blue), HPIV1 (green), and HPIV3 (purple) in HeLa cells. **(D)** Antibodies were also tested for hemagglutination inhibition activity (HAI) against SeV. Antibodies 1A6, 3G12 **(E)** and 11H12 **(F)** were assayed for ability to bind (X-axis) BsrT7 cells infected with SeV containing an eGFP reporter (green, Y-axis) by flow cytometry, as compared to uninfected parental BsrT7 cells (blue) or BsrT7 cells stably and inducibly expressing SeV F (orange).*

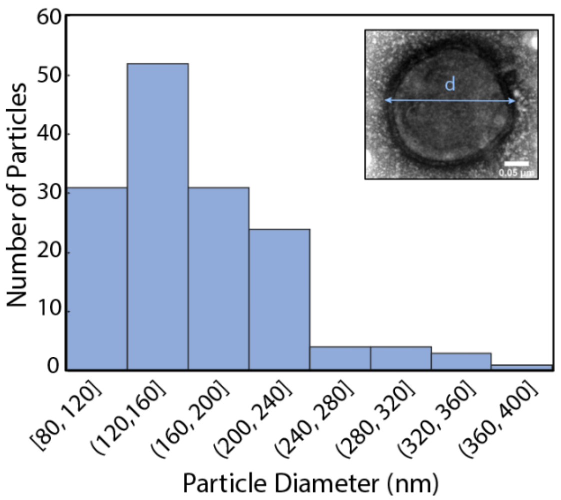

***Figure S10. Distribution of Sendai virus particle sizes as measured by negative stain TEM.*** *Particle outer diameter was measured for 150 SeV particles. The inset image shows an example virion, scale bar = 0.05 µm.*
